# CIViC: A knowledgebase for expert-crowdsourcing the clinical interpretation of variants in cancer

**DOI:** 10.1101/072892

**Authors:** Malachi Griffith, Nicholas C Spies, Kilannin Krysiak, Adam C Coffman, Joshua F McMichael, Benjamin J Ainscough, Damian T Rieke, Arpad M Danos, Lynzey Kujan, Cody A Ramirez, Alex H Wagner, Zachary L Skidmore, Connor J Liu, Martin R Jones, Rachel L Bilski, Robert Lesurf, Erica K Barnell, Nakul M Shah, Melika Bonakdar, Lee Trani, Matthew Matlock, Avinash Ramu, Katie M Campbell, Gregory C Spies, Aaron P Graubert, Karthik Gangavarapu, James M Eldred, David E Larson, Jason R Walker, Benjamin M Good, Chunlei Wu, Andrew I Su, Rodrigo Dienstmann, Steven JM Jones, Ron Bose, David H Spencer, Lukas D Wartman, Richard K Wilson, Elaine R Mardis, Obi L Griffith

## Abstract

CIViC is an expert crowdsourced knowledgebase for Clinical Interpretation of Variants in Cancer (www.civicdb.org) describing the therapeutic, prognostic, and diagnostic relevance of inherited and somatic variants of all types. CIViC is committed to open source code, open access content, public application programming interfaces (APIs), and provenance of supporting evidence to allow for the transparent creation of current and accurate variant interpretations for use in cancer precision medicine.

---

Precision medicine refers to the use of prevention and treatment strategies that are tailored to the unique features of each individual and their disease^1^. In the context of cancer this might, for example, involve the identification of specific genomic mutations shown to predict response to a targeted therapy. The biomedical literature describing these associations is large and growing rapidly. As a result, the interpretation of individual variants observed in patients has become a bottleneck in clinical sequencing workflows^2^. Many cancer hospitals and research centers are engaged in separate efforts to interpret cancer-driving mutations and genes in the context of clinical relevance. These efforts are largely occurring within independent “information silos”, producing interpretations that require constant updates, lack community consensus, and involve intense manual input. Furthermore, the resulting variant interpretations exist largely in private or encumbered databases (**Supplementary Table 1**).

Estimates of the proportion of cancer patients that would benefit from comprehensive molecular profiling vary substantially^3^, in part due to the lack of both a community consensus definition of actionability and a comprehensive catalogue of specific clinical variant interpretations. Achieving the goals of precision medicine will require this information to be centralized, freely accessible, openly debated, and accurately interpreted for application in the clinic. Existing efforts to facilitate clinical interpretation of variants include the Gene Drug Knowledge Database^4^, the Database of Curated Mutations (http://www.docm.info/), ClinVar^5^, ClinGen^6^, PharmGKB^7^, Cancer Driver Log^8^, My Cancer Genome^9^, Jax-Clinical Knowledgebase^10^, the Personalized Cancer Therapy Knowledgebase (https://pct.mdanderson.org/), the Precision Medicine Knowledgebase (https://pmkb.weill.cornell.edu/), the Cancer Genome Interpreter (https://www.cancergenomeinterpreter.org/), OncoKB (http://oncokb.org/), and others (**Supplementary Table 1**). These resources often have barriers to widespread adoption, including some combination of (1) no public access to content, (2) restrictive content licenses, (3) no public application programming interface (API), (4) no bulk data download capabilities, and (5) no mechanism for rapid improvement of the content (see **Supplementary Table 1** for a detailed feature comparison). To address these limitations, we present ‘CIViC’, an open access, open source knowledgebase for expert-crowdsourcing of Clinical Interpretation of Variants in Cancer (www.civicdb.org; **Figure 1**).

**Figure 1.**
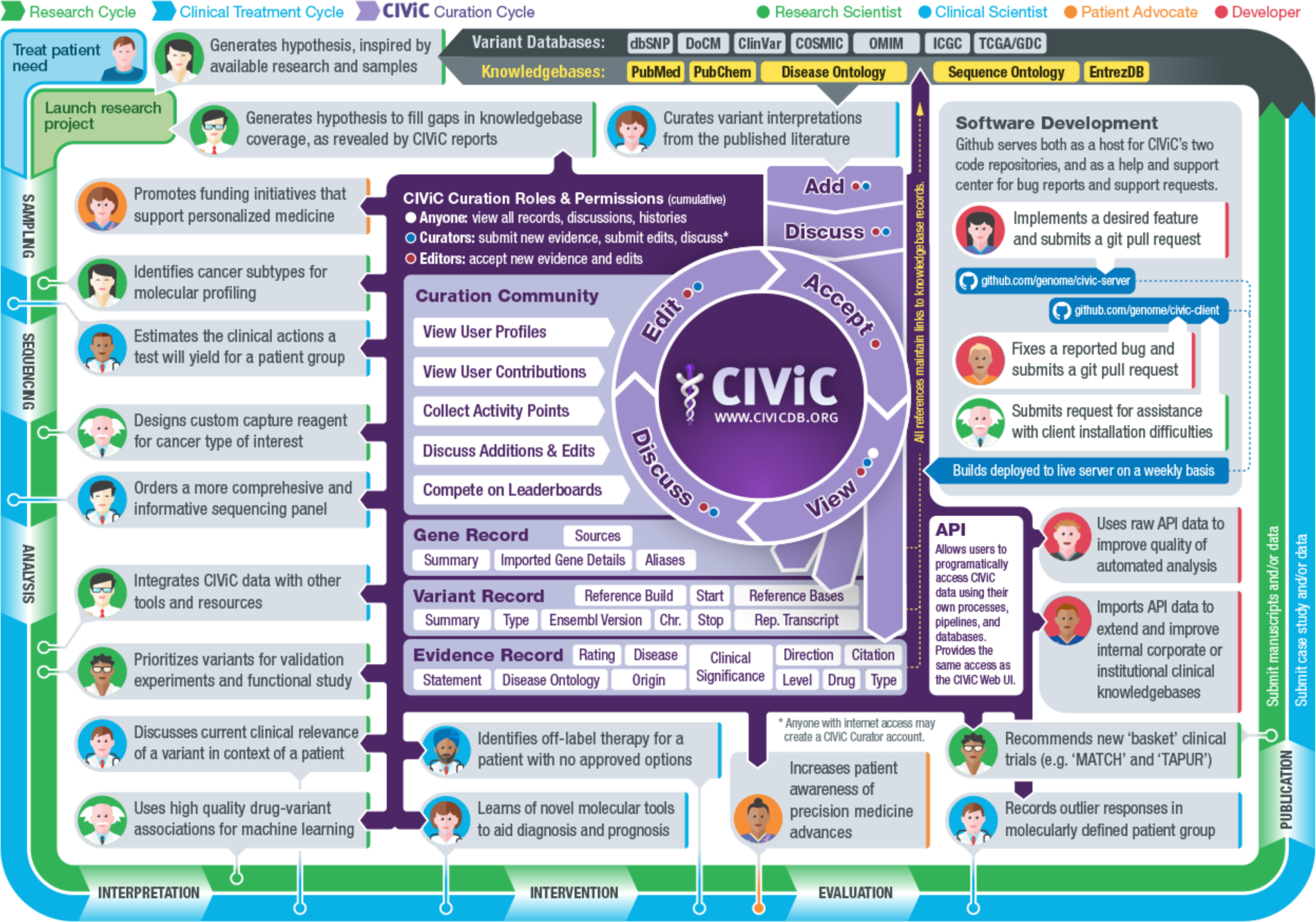
Contribution of CIViC to the precision cancer treatment cycle. The following diagram summarizes how research, clinical treatment, and CIViC knowledge curation are interrelated. The CIViC knowledgebase aims to develop clinical interpretations for raw cancer variant observations stored in large variant databases (grey). Each CIViC variant interpretation is based on published evidence and leverages complementary knowledgebases and ontologies wherever possible (yellow). The precision medicine clinical treatment cycle (blue) and research cycle (green) both involve sampling, sequencing, analysis, interpretation, intervention, evaluation, and publication. These cycles start with hypothesis generation, followed by research projects or clinical trials, and dissemination of their findings. Examples of how each stage specifically relates to or benefits from the CIViC resource are represented by ‘persona’ icons for the four types of CIViC stakeholders: research scientists (green), clinical scientists (blue), patient advocates (orange), and developers (red). Each is accompanied by a brief description of a possible research, clinical, outreach, or software development action. In the center of the diagram, key features of the CIViC interface and data model are summarized (purple). These include the roles and permissions of CIViC users, especially consumers of the content, curators, and editors. Members of the CIViC community participate by adding, editing, discussing, and approving individual evidence records and summaries that support the clinical interpretation of cancer variants. Anyone willing to login may assume the role of curator, but contributions must be reviewed by expert editors prior to acceptance.

The critical distinguishing features of the CIViC initiative, compared to the resources cited above, stem from its strong commitment to openness and transparency. We believe that these principles (see inset **Box 1**) are necessary for widespread adoption of such a resource. The target audience of CIViC is deliberately broad, encompassing researchers, clinicians, and patient advocates. CIViC is designed to encourage development of community consensus by leveraging an interdisciplinary, international team of experts collaborating remotely within a centralized curation interface. Variant interpretations are created with a high degree of transparency and provenance. The interface is designed to help keep interpretations current and comprehensive, and to acknowledge the efforts of creators (**Supplementary Figure 1**). It accepts public knowledge contributions but requires that experts review these submissions.

The manner in which the clinical relevance of variants in cancer is presented in the published literature is highly heterogeneous. In order to represent this data in a more easily interpretable and consistent fashion, the CIViC data model is highly structured and ontology-driven (**Supplementary Figure 2**). Clinical interpretations are captured and displayed as evidence records consisting of a free-form ‘evidence statement’ and several structured attributes. Each evidence record is associated with a specific gene, variant, disease, and clinical action. Evidence records belong to one of three evidence types indicating whether a variant is predictive of response to therapy, prognostic, and/or diagnostic. Evidence records are also assigned to an evidence level ranging from established clinical utility (level A) to preclinical (level D) or inferential (level E) evidence ( **Supplementary Figure 3** and **Supplementary Figure 4**) and the quality of the underlying published evidence is rated from one to five stars. As evidence records accumulate for a single variant, they are, in turn, synthesized into a human-readable ‘variant summary’ of the variant’s overall significance in cancer. Variants can also be aggregated into ‘variant groups’ that share a clinical significance (e.g. ‘imatinib resistance’). All variant types are supported (including structural variants, RNA fusions, and other expression events) as well as all variant origins (somatic, germline mutation, and germline polymorphism). Genomic coordinates, transcript identifiers, and variant synonyms are determined by curators, reviewed by editors, and stored in a standardized format for each variant. Each variant is associated with a single gene, and each gene provides a ‘gene summary’ synthesizing all of the variants it contains. Additional gene information is imported through the MyGene.info annotation API^11^, allowing users to focus curation effort on clinical impact, and not repeat the efforts of other databases. Integration of public ontologies and databases, such as the Disease Ontology^12^, the Sequence Ontology^13^, and PubChem^14^, allow CIViC’s data to be formally structured (**Supplementary Figure 5**) and integrated with other resources. This structure provides computationally accessible information integrated with human interpretable content and the flexibility to capture key details unique to the plethora of variants and experiment types being interpreted (refer to **Supplementary Methods** for implementation details).

CIViC currently contains 1,411 curated interpretations of clinical relevance for 560 variants affecting 235 genes (**Supplementary Figure 6**). These interpretations were curated from 918 published studies by 37 CIViC curators. CIViC evidence records are currently biased towards somatic alterations, supporting positive associations with treatment response and supported by a wide range of evidence levels and trust ratings (**Supplementary Figure 7**). At least one evidence records has been created for 144 cancer subtypes and 271 drugs, with some bias towards lung, breast, leukemia, colorectal and skin cancer, and associated targeted therapies (**Supplementary Figure 8**). Supporting publications for these interpretations come from a large number of journals, primarily over the last five years, and tend to provide just one or two evidence records each (**Supplementary Figure 9**). From June 2015 to April 2016, external curators (not affiliated with Washington University) contributed 41.6% of the evidence statements within the knowledgebase (**Supplementary Figure 6B**). To date, submissions, revisions, comments, and expert reviews have produced 8,143 distinct curation actions. These numbers continue to grow. More than 10,000 users have accessed CIViC interpretations from academic, governmental, and commercial institutions around the world (**Supplementary Figure 6C-D**). Early adopters of CIViC include leaders in developing cancer genomics pipelines^15^ and the UCSC Genome Browser^16^. Early curation/content partners include the Gene Drug Knowledgebase^4^, and the Personalized Oncogenomics Program^17^. The CIViC resource is freely accessible without login and no fees or exclusive access will be introduced in the future. Both academic and commercial adoption are free and encouraged. The variant and gene summaries, with additional statistics summarizing the level of supporting evidence in CIViC, can be automatically incorporated into clinical reports using the CIViC API or bulk data releases (updated nightly, with stable monthly releases) (**Figure 1**). The source code for the CIViC website and public API are released under an open source license (MIT), and all curated content within CIViC is released under an open access license (Creative Commons Public Domain Dedication, CC0). The unencumbered availability of the CIViC bulk data releases, lack of requirements to establish a licensing agreement, well-documented public API, and use of a structured data model and ontologies, encourages adoption of CIViC in clinical workflows. As the user base grows, the number of experts with a vested interest in the content will increase, driving community engagement and increasing curation from external users.

A critical emphasis in the growth of CIViC content will be in maintaining high quality. The curation workflow of CIViC (**Supplementary Figure 10**) requires agreement between at least two independent contributors prior to acceptance of new evidence or revisions of existing content (**Supplementary Figure 11** and **Supplementary Figure 12**). At least one of these users must be an expert editor, and editors are barred from approving their own contributions. CIViC includes features such as typeahead suggestions, automatic warning of possible duplicates, detailed documentation in all entry forms, and input validation, to encourage high quality data entry. To facilitate team curation efforts (**Supplementary Figure 10**), the CIViC interface also includes features such as subscriptions, notifications, mentions, and ‘macros’ that assist communication between curators and editors (e.g. macro assist in linking a discussion to related evidence records, revisions, etc.). Curators can also use an advanced search interface to generate and share complex queries of CIViC data (currently 40 features of evidence, variant, and gene records) that help guide curation effort (**Supplementary Figure 13**). Many of these features were inspired by the ‘best practices’ of active online collaborative research and software development platforms including BioStars^18^ and GitHub [https://github.com/].

A major challenge to the success of CIViC is the scope and complexity of the knowledge that needs to be summarized. The American College of Medical Genetics and Genomics (ACMG) and the Association for Molecular Pathology (AMP) recently reported on the variability between nine labs in clinical interpretations of variants relevant to Mendelian diseases^19^, a field where the ACMG-AMP have proposed detailed Standards and Guidelines for variant classification^20^. This report identified a low rate of interpretation agreement between labs (34% concordance). However, discussion and criteria review were able to more than double this concordance, further demonstrating the need for and success of open discourse in clinical variant interpretation^19^. At present, there is remarkably low overlap between the publications cited for clinical interpretation of cancer variants by multiple efforts that nominally share the same goals (1.6% - 71.6% overlap but generally less than 25%; **Supplementary Table 2**). This suggests that no single effort has comprehensively identified or summarized even the most relevant literature in this area, further illustrating the high curation burden involved. Conversely, these small overlaps emphasize the importance of reducing duplication of effort moving forward, especially considering the vastness of the existing literature and its tremendous growth rate. In CIViC, curation efforts to date have focused on variants relevant to cancer types of particular focus at our center (e.g. acute myeloid leukemia, breast, and lung; **Supplementary Figure 8**), on variants identified as high priority by early CIViC partners^4,17^, and on variants targeted by proof-of-principle precision medicine “basket” clinical trials such as the NCI-MATCH (also known as EAY131 or NCT02465060). Our ability to provide expertise in these areas is complemented by the expert knowledge of other groups and organizations, making CIViC a more well-rounded, comprehensive resource that would not be possible with a “siloed data” approach.

CIViC is uniquely designed to address the immense challenge of open discussion and review of variant interpretations. To our knowledge, CIViC is the only variant interpretation effort currently capable of leveraging community experts, and additionally has the most open model (open access content, open source code, and open API). We believe that this open strategy represents a sustainable model for achieving current, standardized, and comprehensive interpretations of the clinical relevance of cancer variants. As the community of contributors grows, an increased incentive will emerge to help keep CIViC updated with cutting edge clinical trial and FDA IND findings. Since we have created a comprehensive and modern application programming interface (API), centers can rapidly integrate CIViC into automated clinical report generation for gene panel, exome, whole genome, and RNA sequencing of tumor samples. We invite all researchers and clinicians engaged in clinical interpretation of variants to join the community at CIViC (www.civicdb.org).

## Author Contributions

MG, NCS, KK, and OLG wrote the paper with input from RL, ERM, AHW, ACC, JFM, BJA, CAR, CJL, DTR and GCS. ACC led the back end code development with contributions from MG, NCS, JFM, OLG, KK, GCS, and GK. JFM led the front end code development with contributions from APG, ACC, KK, MG, OLG, and NCS. BMG, AIS, and SJMJ contributed ideas relating to crowdsourcing functionality. JME, DEL, and JRW contributed software engineering expertise. Substantial curation efforts were contributed by MG, NCS, KK, BJA, DTR, AMD, LK, CAR, AHW, ZLS, CJL, MJ, RLB, RL, EKB, NMS, MB, LT, MM, AR, KMC, RB, DHS, LDW, ERM and OLG. Guidance on developing CIViC for clinical applications was provided by RB, DHS, and LDW. Trainee supervision and project leadership were provided by MG, RKW, ERM, and OLG.

## Acknowledgements and Funding

First and foremost we are grateful to the community of curators, editors, domain experts, and users who make CIViC possible. This work was supported by a grant to Richard K. Wilson from the National Human Genome Research Institute (NHGRI) of the National Institutes of Health (NIH) under Award Number U54HG003079. DTR was supported by the German Federal Ministry of Education and Research under Award Number 031L0030E and 031L0023B. MG is supported by the NHGRI under Award Number K99HG007940. OLG is supported by the National Cancer Institute (NCI) of the NIH under Award Number K22CA188163. LDW is supported by the NCI under Award Number K08CA166229. The CIViC project is also supported by the NCI under Award Number U01CA209936 to OLG (with MG and ERM as co-PIs). The content of this manuscript is solely the responsibility of the authors and does not necessarily represent the official views of the NIH.

### Box 1. CIViC principles

1. **Interdisciplinary**. An interdisciplinary approach is needed to combine the expertise of genome scientists, health care providers, patient advocates, and others.
2. 2. **Community consensus.** The interpretations of clinical actionability required to enable precision medicine should be freely available and openly discussed across a diverse community. To facilitate consensus building, the interface must support direct contribution from members of the community.
3. **Transparency.** Content should be created with transparency, kept current, be comprehensive, track provenance, and acknowledge the efforts of its creators.
4. **Computationally accessible.** The interface should be both structured enough to allow computational data mining (via APIs) and agile enough to handle the product of openly-debated human interpretation.
5. **Freely accessible.** Curated knowledge will remain free and can be accessed anonymously without login unless the user wishes to contribute to content. No fees will be introduced.
6. **Open license.** CIViC will encourage both academic and commercial engagement through flexible licensing. Access will not be restricted by exclusive licensing.

